# Deep learning to extract the meteorological by-catch of wildlife cameras

**DOI:** 10.1101/2023.09.25.558780

**Authors:** Jamie Alison, Stephanie Payne, Jake M. Alexander, Anne D. Bjorkman, Vincent Ralph Clark, Onalenna Gwate, Maria Huntsaar, Evelin Iseli, Jonathan Lenoir, Hjalte Mads Rosenstand Mann, Sandy-Lynn Steenhuisen, Toke Thomas Høye

## Abstract

Microclimate – proximal climatic variation at scales of meters and minutes – can exacerbate or mitigate the impacts of climate change on biodiversity. However, most microclimate studies are temperature-centric, and do not consider meteorological factors such as sunshine, hail and snow. Meanwhile, remote cameras have become a primary tool to monitor wild plants and animals, even at micro-scales, with deep learning tools to rapidly convert images into ecological data. However, deep learning applications for wildlife imagery have focused exclusively on living subjects. Here, we identify an overlooked opportunity to extract latent, yet ecologically relevant, meteorological information.

We produce an annotated image dataset of micrometeorological conditions across 49 wildlife cameras in South Africa’s Maloti-Drakensberg and the Swiss Alps. We train ensemble deep learning models to classify conditions as overcast, sunshine, hail or snow. Our best model achieves 91.7% accuracy on test cameras not seen during training. Furthermore, we show how effective accuracy is raised to 96% by disregarding 14.1% of classifications where ensemble member models did not reach a consensus. Unlike previous studies we test model performance in remote and novel locations, distinguishing overcast and sunny conditions in Svalbard, Norway with 79.3% accuracy (93.9% consensus accuracy).

Our model rapidly classifies sunshine, snow and hail in almost 2 million unlabeled images. Resulting micrometeorological data illustrated common seasonal patterns of summer hailstorms and autumn snowfalls across mountains in the northern and southern hemispheres. However, daily patterns of sunshine and shade diverged between sites, impacting daily temperature cycles. Crucially, we leverage micrometeorological data to demonstrate that (1) experimental warming using open-top chambers shortens early snow events in autumn, and (2) image-derived sunshine marginally outperforms sensor-derived temperature when predicting bumblebee foraging. These methods generate novel micrometeorological variables in synchrony with biological recordings, enabling new insights from an increasingly global network of wildlife cameras.

## Introduction

Climate change is redistributing species in space and time, with profound impacts on ecosystem function and human wellbeing (Pecl et al., 2017). While biodiversity impacts of climate change are usually reported at coarse spatial resolutions across large spatial extents (Lenoir et al., 2020), species often respond to climate at much finer scales (Lembrechts et al., 2019; Maclean & Early, 2023; Potter et al., 2013). Furthermore, studies of the biodiversity impacts of climate change are traditionally temperature-centric (Antão et al., 2020). As the study of microclimate has expanded, this temperature-centricity has remained (Maclean et al., 2021), due in part to the availability of inexpensive and easy-to-use temperature loggers (Lembrechts et al., 2020). Still, non-temperature aspects of climate and meteorology have profound, fine-scale impacts on species and communities. Solar radiation (i.e., sunshine) underpins ambient temperature, but also directly impacts plant growth and animal behavior (Möhl et al., 2020; Roales et al., 2013; Valladares et al., 2016).

Furthermore, snow cover severely constrains the onset of plants’ growing and flowering seasons (Möhl et al., 2022), impacting plant-pollinator interactions (Gillespie et al., 2016; Gillespie & Cooper, 2022). Hail receives very little attention in ecological research, yet presents a direct physical threat that can devastate poorly-adapted plant species (Fernandes et al., 2011). As the frequency of extreme weather events increases (IPCC, 2023), it is vital that sensor networks capture not only temperature, but also a range of other fine-scale meteorological variables.

Meanwhile, novel technology is revolutionizing the monitoring of ecological communities, generating data with unprecedented temporal continuity and resolution (August et al., 2015; Besson et al., 2022). Wildlife cameras – *in situ* autonomous cameras that record wildlife – are particularly promising. Camera traps are an established method to monitor large animals (Burton et al., 2015), but wildlife cameras are now regularly deployed to study small mammals and birds (Ortmann & Johnson, 2021), insects and other arthropods (Høye et al., 2021), and plants (Katal et al., 2022). For vegetation, the “PhenoCam” approach has gained traction, capturing community-level characteristics such as plant productivity (Brown et al., 2016). However, the volume of images from wildlife cameras has proven difficult to manage, so algorithms are being developed to automatically extract ecological data (Høye et al., 2021; Tuia et al., 2022). Deep learning models are a prevalent family of algorithms used to detect, classify and track animals in images (Norouzzadeh et al., 2018). For plants, algorithms can be trained to flag phenological events such as the onset of budburst or flowering (Katal et al., 2022), or to detect and track individual flowers (Mann et al., 2022).

The consistency of image-based monitoring allows for incidental recording of non-target biota, known as “by-catch”. For example, a camera trap network intended to study the Eurasian lynx in the Swiss Jura mountains has proven useful to explore habitat use by wild boar and roe deer (Wevers et al., 2021). However, it is increasingly recognized that wildlife cameras also capture non-target abiotic information, including meteorological conditions not easily captured with alternative sensors (Hofmeester et al., 2020). Furthermore, capturing both biotic and abiotic data with a single sensor ensures that they are measured simultaneously, at equivalent spatial and temporal scales. Several studies have manually extracted the presence, cover or depth of snow in ecological imagery (Lumbrazo et al., 2022). Furthermore, some studies automate the quantification of snow in PhenoCam images (Caparó Bellido & Rundquist, 2021; Julitta et al., 2014), or the classification of frost (Noh et al., 2021). However, the use of a single, efficient classifier to simultaneously detect sunshine and frozen precipitation in wildlife images has not been explored.

Working with wildlife imagery from matching experimental sites in mountains in the northern (Switzerland; CH) and southern (South Africa; ZA) hemispheres, we train ensemble deep learning models to detect micrometeorological conditions of sunshine, hail, and snow. Our objectives are as follows: (1) Build a dataset and train deep learning classifiers to derive micrometeorological conditions in images; (2) evaluate different data-splitting and ensemble approaches to maximize model performance with out-of-sample and out-of-distribution test datasets; and (3) demonstrate the application of image-derived micrometeorological variables for ecological research. For this last objective, we classify conditions in almost 2 million unlabeled images in CH and ZA. Then we (i) use snow in ZA images to determine the effects of experimental warming on snowmelt, and (ii) use sunshine in CH images to explore the relative importance of sunshine and ambient temperature for bumblebees foraging at high elevations. Our approach efficiently attaches micrometeorological data to remote biodiversity observations. This is particularly useful for biodiversity monitoring in mountains, which are characterised by high levels of endemism, complex topography, and microclimatic heterogeneity (Spehn & Körner, 2005).

## Methods

### Study area

Imaging was carried out over 49 1×1m montane grassland plots across four replicated experimental field sites. In the Calanda massif in the Alps in Switzerland (CH), 24 plots were imaged from June 2021 to October 2021, of which eight were located at a low elevation site (46.869266° N 9.490098° E; 1,440m elevation) and 12 were located at a high elevation site (46.887824° N 9.489510° E; 2,000m). In the Sentinel region of the Maloti-Drakensberg in South Africa (ZA), 25 subplots were imaged from November 2021 to April 2022, of which five were located at a low elevation site (28.679532° S 28.894816° E; 2,200m) and 20 were located at a high elevation site (28.754951° S 28.866981° E; 3,060m). The grassland plots we imaged comprised native vegetation, but half of the plots additionally had small plant specimens, of species typical of lower elevations, introduced within them. Half of the high site plots in ZA were also exposed to open top chamber (OTC) warming of approximately 2°C.

### Sensor deployment

A Wingscapes TimelapseCam Pro camera (with LED flash) was mounted on a steel frame over each plot, ca. 62cm from the ground, pointing towards the ground. Each camera captured an area of ca. 30×17cm. All cameras were ‘continuous’, recording images day and night at 5-minute intervals, except for 16 ‘focused’ cameras in CH which recorded at 1-minute intervals between 12.00–15.00 and 01.00–03.00 (Alison et al., 2022). Each camera required eight AA lithium batteries and a 128GB SD card, replaced approximately every two months.

All cameras were equipped with on-board temperature sensors, recording temperature at ca. 62cm above ground with every photograph (either at 5- or 1-minute intervals depending on the camera). At the high elevation sites, we also deployed TMS4 microclimate loggers to record temperature every 15 minutes at -8 cm, 0 cm and 15 cm above ground. We deployed loggers in blocks of six, with three blocks at the CH high site and one block at the ZA high site. Each block included two loggers in vegetated plots with OTC warming, two in vegetated plots without warming, and one each in bare soil plots with and without OTC warming. Loggers were not targeted to the same plots as cameras, though some of the same plots were sampled by coincidence.

### Image labelling

We sorted a total of 8,953 images into four classes based on micrometeorological conditions: overcast, sunshine, hail or snow (Fig. 1; further details in Appendix S1). First, to generate training data representing the entire recording period, we labelled a systematic sample of 6,205 images. We sampled one image per hour in both ZA and CH, cycling through the continuous cameras. In CH, we included an additional one image per hour cycling through all focused cameras (five hours per day). The systematic sample yielded 1,320 sunshine, 110 snow, 39 hail, and 4,736 overcast images. Second, to generate adequate training data and improve detection of snow and hail, we took a strategic sample of 2,748 images that was informed by the systematic sample. The strategic sampling protocol had three tiers, such that the rarest meteorological events were sampled more intensively (i.e. oversampled; Table S1). The strategic sample yielded an additional set of 266 sunshine, 923 snow, 803 hail and 756 overcast images. CH images were labelled by JA, and ZA images by SP, using VGG Image Annotator (VIA v2.0.11; https://www.robots.ox.ac.uk/~vgg/software/via/).

**Figure 1.**
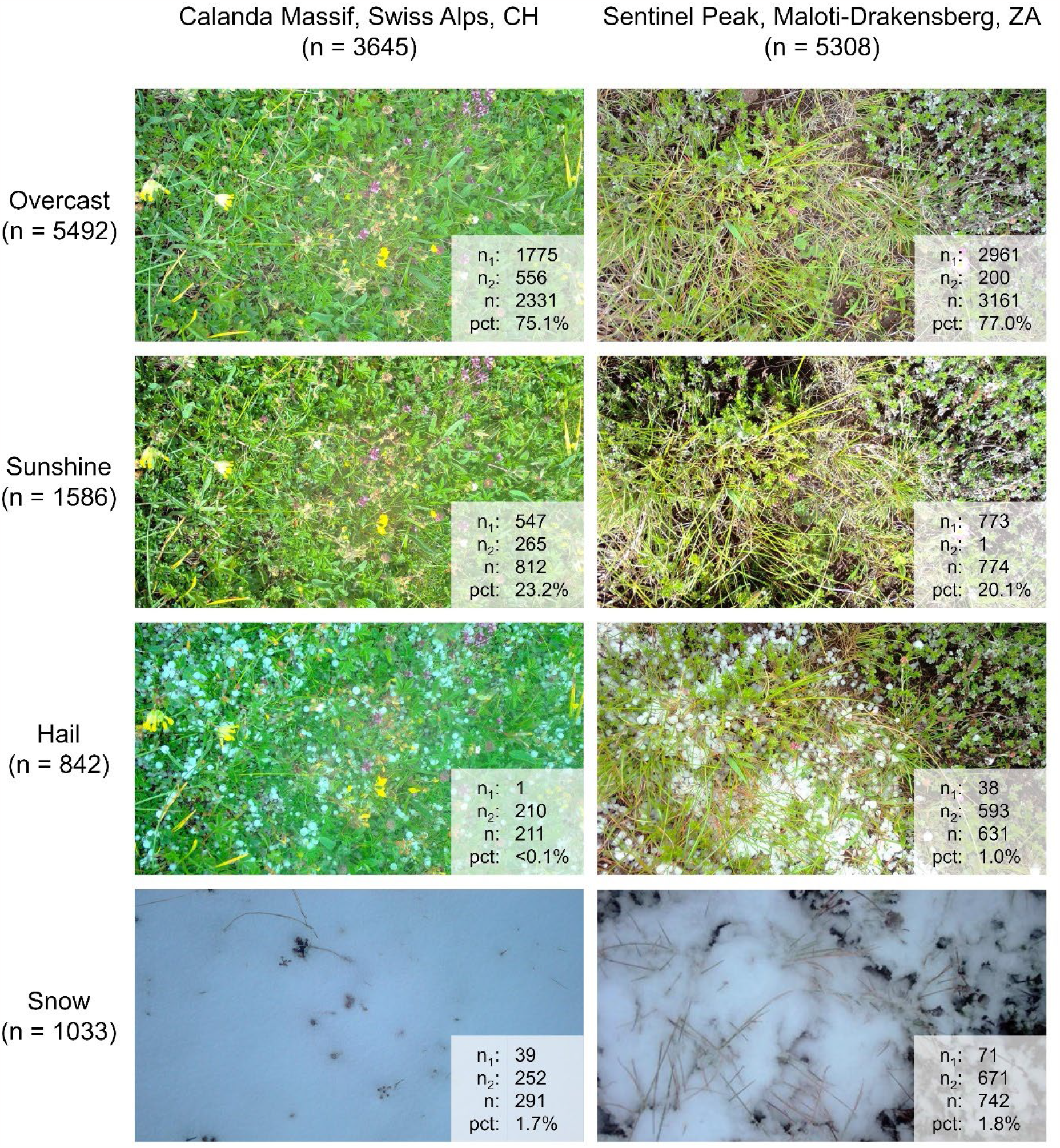
Examples of the four classes of micrometeorological conditions in two representative high elevation plots in Switzerland (CH) and South Africa (ZA). Total numbers of labelled images (n) are shown across classes and regions (including high and low elevation sites). The breakdown of labels across the systematic sample (n_1_) and the strategic sample (n_2_) is also shown, as is the representation of each condition within the systematic sample for a given region (pct).

### Model training

We trained convolutional neural networks (CNNs) to classify conditions in images as overcast, sunshine, hail or snow using the *Keras* python library (Chollet, 2015). Specifically, we fine-tuned the MobileNet network (Howard et al., 2017) pretrained on ImageNet (Russakovsky et al., 2015), representing a lightweight and efficient family of CNNs that has been shown to perform well for image classification in ecology (Besson et al., 2022; Schneider et al., 2022). To adapt the model to predict just four classes, we removed the top layer of the network and added a custom softmax activation layer with a flattened input. Training images were rescaled to 224×224px and put through an augmentation pipeline to reduce overfitting. Augmentation involved random vertical or horizontal flipping and up to 45° random rotation in any direction. Following preliminary tests, we trained the entire network in two stages using the Adam optimizer and a batch size of 128. First, we trained for five epochs specifying a learning rate of 0.001, to bring models rapidly towards a solution. Second, we trained the entire network for up to 200 epochs, specifying a learning rate of 0.0001. The change in learning rate allowed models to reach an optimal solution while minimizing spurious fluctuations in validation loss. Appendix S2 gives a primer on CNN parameters and concepts.

### Model validation

We used model validation with early stopping to minimize overfitting to the training data. We stopped training if validation performance did not improve for 30 epochs and saved the model from the epoch with the best validation performance (concepts explained in Appendix S2). To account for variance related to data splitting for validation, we used cross-validation. Specifically, we carried out a 5-fold cross-validation in which each of five mutually exclusive data folds is iteratively treated as the validation dataset (e.g. Fig. S1, inner validation loop). Additionally, we compared two data splitting methods: “Cis” and “Trans” (Beery et al., 2018, 2020). Cis validation involves random splitting of images from all cameras. Trans validation splits cameras rather than images, such that images from one camera are always constrained to a single fold (Fig. S1). With the Cis validation method, the effective size of the training set is larger, because more camera positions are seen during training. However, the Trans validation method ensures that images from the same camera are not represented in both training and validation data. For both Cis and Trans validation, folds were stratified with respect to the four study sites (Wieczorek et al., 2022).

### Model aggregation

We produced ensembles of sets of five “member models”, representing the five iterations of each 5-fold validation loop, in a procedure called cross-validation aggregation (crogging; Barrow & Crone, 2013; see Appendix S3 for further details). The crogging procedure produces ensemble models in which each observation contributes to both model training (in all but one member model) and model validation by early stopping (in one member model). Averaging across models which use distinct data folds for validation can improve generalization performance and model stability (Barrow & Crone, 2016; Krogh & Vedelsby, 1994). Ensemble models were produced under both Cis and Trans validation methods. For each 5-fold validation loop, five Cis member models and five Trans member models were aggregated into ensembles by unweighted averaging of output softmax probabilities.

### Model testing and deployment

We aimed for models that would transfer to (1) novel camera positions within our sites and (2) novel sites with a similar recording protocol. To test transferability to novel positions within our sites, each 5-fold validation loop was nested within an outer 6-fold test loop (e.g. Fig. S1). In other words, we used a nested cross-validation or “double-cross” to obtain a robust and unbiased measure of model generalization (Cawley & Talbot, 2010). We randomly split 49 cameras into six folds that were stratified across the four sites (Wieczorek et al., 2022). During each test loop iteration, one of six folds was withheld as a test dataset. We compared model accuracy and macro-F1 (the mean of class F1 scores) following Eq. 1-4 where: *TP* = true positives; *TN* = true negatives; *FP* = false positives; *FN* = false negatives; *Pr* = class precision; and *Re* = class recall. Class F1 scores represent class-level performance, and macro-F1 represents overall model performance while accounting for class-imbalance.

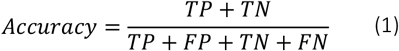

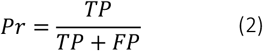

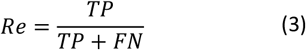

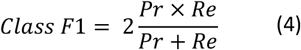

To test model transferability to novel sites, we trained three production models by incorporating the test data for training and validation. Specifically, we trained a six-fold Trans ensemble and a six-fold Cis ensemble. Novel site performance was then tested using images from Svalbard, Norway. In a separate study of pollination of *Silene acaulis*, six plots in Bjørndalen (78.21660° N 15.33280° E; 40m elevation) were imaged at 1-minute intervals in June and July 2020 using the same imaging methods. However, some images were cropped slightly to zoom in on *S. acaulis* during annotation. The first 11 images of every hour of every day were labelled for sunshine (n = 957) and overcast (n = 6222) conditions by MH.

Finally, we deployed the Trans ensemble model to predict micrometeorological conditions across the full set of 1,934,044 images taken across all four sites in CH and ZA. We validated our time-series of micrometeorological conditions against temperature data from TMS4 loggers (Wild et al., 2019). We then analyzed micrometeorological conditions to explore (1) effects of OTC warmed and unwarmed treatments on snow cover in ZA, and (2) effects of sensor-derived temperature and image-derived sunshine on *Bombus* visitation to *Trifolium pratense* in CH (using data previously published by Alison et al., 2022). For (2), we calculated the daytime degree days recorded by on-board temperature sensors (sum of mean daily temperatures above 0°C) and sunshine days (sum of mean daily proportional sunshine) during flowering of each inflorescence. We compared AIC of linear models predicting ln(number of visits) of each inflorescence. To further visualize the niche preferences of *Bombus* bees, we calculated mean temperature and proportional sunshine within an hour either side of each *Bombus* foraging event. This was overlaid on the distribution of temperature and sunshine during daytime hours in which 32 *T. pratense* inflorescences were flowering.

## Results & Discussion

Wildlife cameras capture details about species’ abiotic environments, and not just their behaviors, life cycles, and interactions (Hofmeester et al., 2020). We find that conditions such as sunshine, snow and hail can be readily and automatically extracted from wildlife imagery. Furthermore, we show how such micrometeorological data are complementary to sensor-derived temperature data.

### Model performance

Ensemble models were highly transferable to novel camera positions, consistently achieving higher accuracy and macro-F1 than member models (Table 1). Ensemble models also outperformed full models, which used all data for training and none for validation, by around 1%. Beyond performance benefits, member models within a given ensemble disagree on 13-14% of predictions (Table 1) and this disagreement forms a useful measure of uncertainty; by omitting non-consensus predictions as uncertain, we raised the effective accuracy of our most accurate ensemble to 96%. Differences between models trained using the Cis and Trans validation methods were negligible, while cross-validation revealed considerable variation when models of the same type were trained and tested on different folds of the data (Table 1, see Appendix S3 for further discussion on validation approaches and ensembling). Our models classified overcast and sunshine conditions more successfully than snow or hail (Fig. 2, Fig. S2), perhaps reflecting the oversampling of snow and hail events. Our ensemble model based on a Trans validation method misclassified 29% of hail images as overcast, 12% of sunshine as overcast, and 3% of overcast as sunshine (Fig. 2).

**Table 1.** Mean (±SD) performance of full models (n=6), ensemble models (n=6) and member models (n=5×6=30) in classification of micrometeorological conditions in out-of-sample test images. Performance metrics include macro-F1, accuracy, and consensus accuracy for ensemble models. Ensemble models are compared to member models under two validation data split methods: “Cis” and “Trans” (see Fig. S1). The consensus accuracy of ensemble models is the accuracy of predictions on which all five member models agree. Rate of consensus is the percentage of test images for which all five member models agree.

| Validation split method | Model type | Macro-F1 (%) | Accuracy (%) | Consensus accuracy (%) | Rate of consensus (%) |
| --- | --- | --- | --- | --- | --- |
| Cis | Ensemble | **89.25** (2.02) | 91.63 (1.96) | 95.89 (1.69) | 87.13 (1.92) |
|  | Member | 88.02 (1.59) | 90.72 (1.67) |  |  |
| Trans | Ensemble | 88.82 (3.33) | **91.70** (2.02) | **96.00** (1.55) | 85.90 (3.33) |
|  | Member | 87.22 (3.02) | 90.41 (2.13) |  |  |
| None | Full | 88.00 (2.61) | 90.94 (2.21) |  |  |

**Figure 2.**
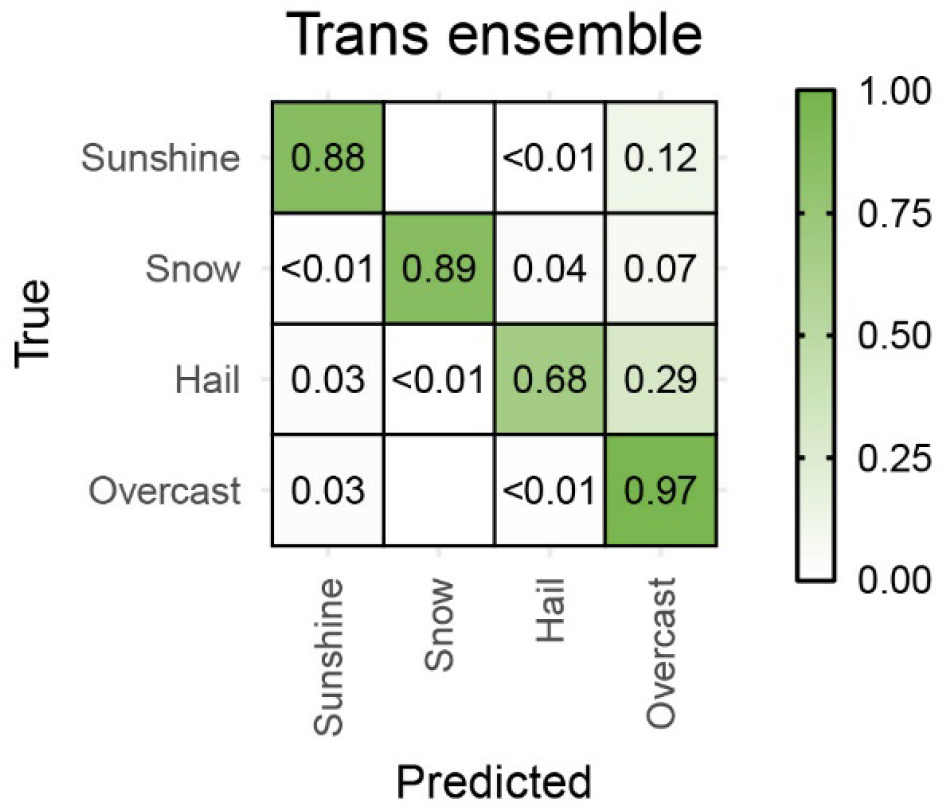
Confusion matrix of predicted micrometeorological conditions across 8,953 images. Six ensemble deep learning models were used to classify images in six distinct folds of hold-out test cameras in a cross-validation process. Data splitting for validation of these models was done using the Trans validation method (see text and Fig. S1 for explanation).

The real-world utility of deep learning models hinges on generalization performance, including transferability to novel sites and conditions. When classifying images of *S. acaulis* across six cameras from a totally independent site in Svalbard, Norway, ensemble models using Cis and Trans validation methods distinguished sunshine from overcast conditions with 79.3% and 72.8% accuracy, respectively (model F1: 70.9% and 65.0%). Furthermore, omitting non-consensus predictions as uncertain, we raised the effective accuracy to 94% and 88.4%, respectively. This predictive performance is impressive given that these out-of-distribution images came from a site >1,000km away, at >1,000m lower elevation, with different height and width compared to training images. For detection of sunshine in images, our models show clear potential to generalize to novel sites with a similar recording protocol.

### Extensive prediction of micrometeorological data

We predicted micrometeorological conditions in almost two million images across CH and ZA sites. A consensus prediction emerged for 87.7% of images, and these predictions were averaged across cameras over time to generate seasonal (Fig. 3) and diel (Fig. 4) micrometeorological profiles. Seasonal profiles revealed a common seasonal pattern of summer hailstorms and autumn snow across mountains in the northern and southern hemispheres (Fig. 3). As expected, during the day there was a striking match between image-derived sunshine and sensor-derived temperatures (Fig. 3). However, nighttime temperatures also appeared warmer following periods of high daytime cloud cover (i.e., low sunshine before nightfall; Fig. 3), as expected under longwave cloud forcing (Ramanathan et al., 1989). Furthermore, diel sunshine profiles revealed that the ZA high elevation site was characterized by morning sunshine and afternoon shade, with subtle impacts on the diel temperature profile (Fig. 4). Such insights would be difficult to obtain without using the meteorological by-catch of wildlife cameras. Crucially, our models extracted the meteorological by-catch of images very rapidly. On a laptop computer, a member model classified two million images in around 20 hours on an Intel Xeon E5-2697A v4 processor with 8 CPUs @∼2.6GHz. Much faster times would be expected if using a GPU.

**Figure 3.**
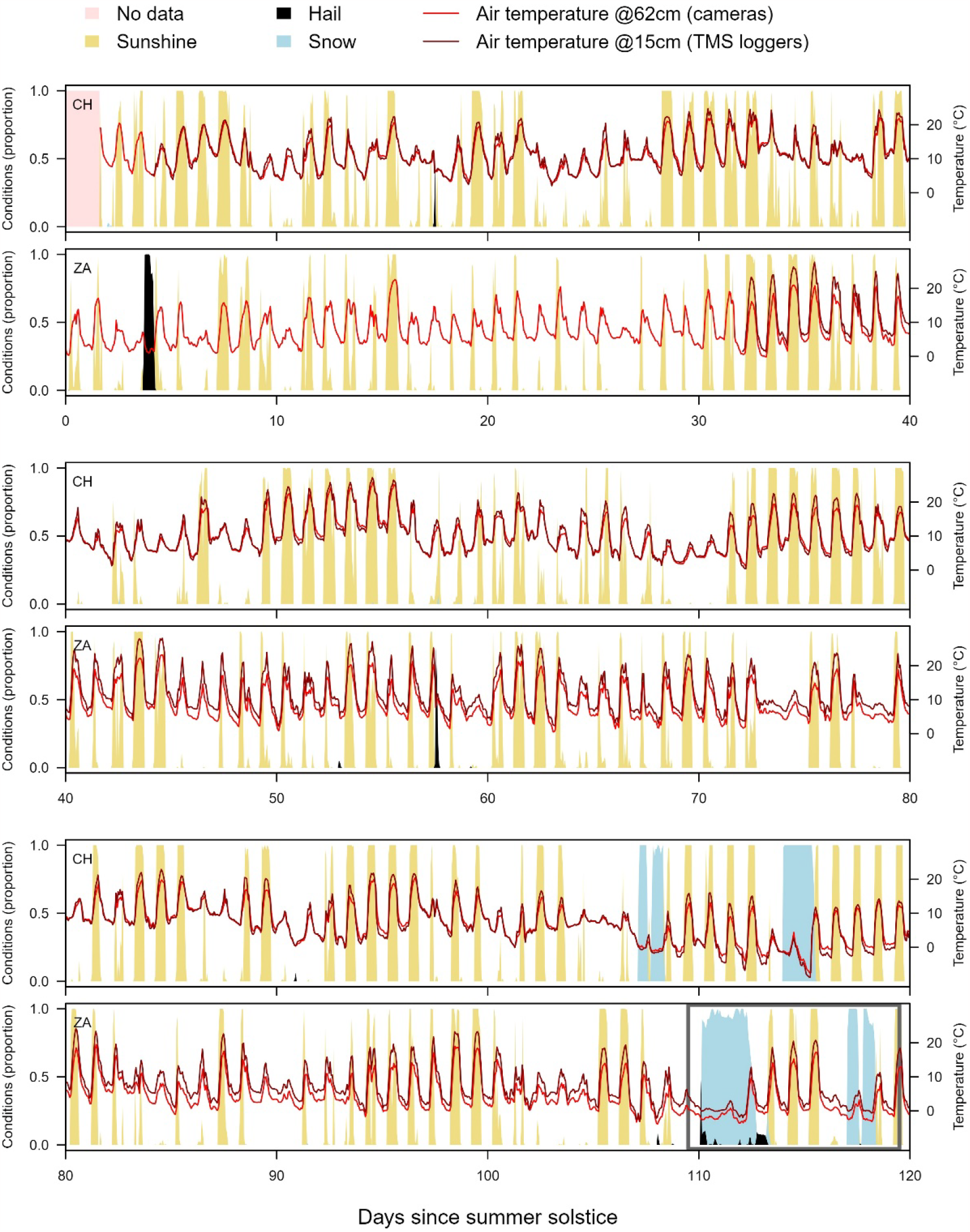
Sunshine (yellow), hail (black) and snow (blue) detected in images throughout summer solstice (top pair), early autumn (middle pair) and late autumn (bottom pair) across two mountain sites. Distinct time series from the northern hemisphere (top of each pair) and the southern hemisphere (bottom of each pair) are aligned based on summer solstice dates of 21^st^ June 2021 in Switzerland (CH) and 21^st^ December in South Africa (ZA). Conditions were classified in all images from continuous cameras at the two high-elevation sites, using a deep learning ensemble trained using the full set of 8,953 labelled images. Data are displayed with a resolution of 1.2 hours (20 time slices per day). We also present mean air temperatures from on-board sensors on cameras, measuring air temperature every 5 minutes at 62cm above ground (red lines), and TMS4 loggers, measuring air temperature every 15 minutes at 15cm above ground (dark red lines). Data in the grey box (bottom-right) are more closely explored in Figure 5.

**Figure 4.**
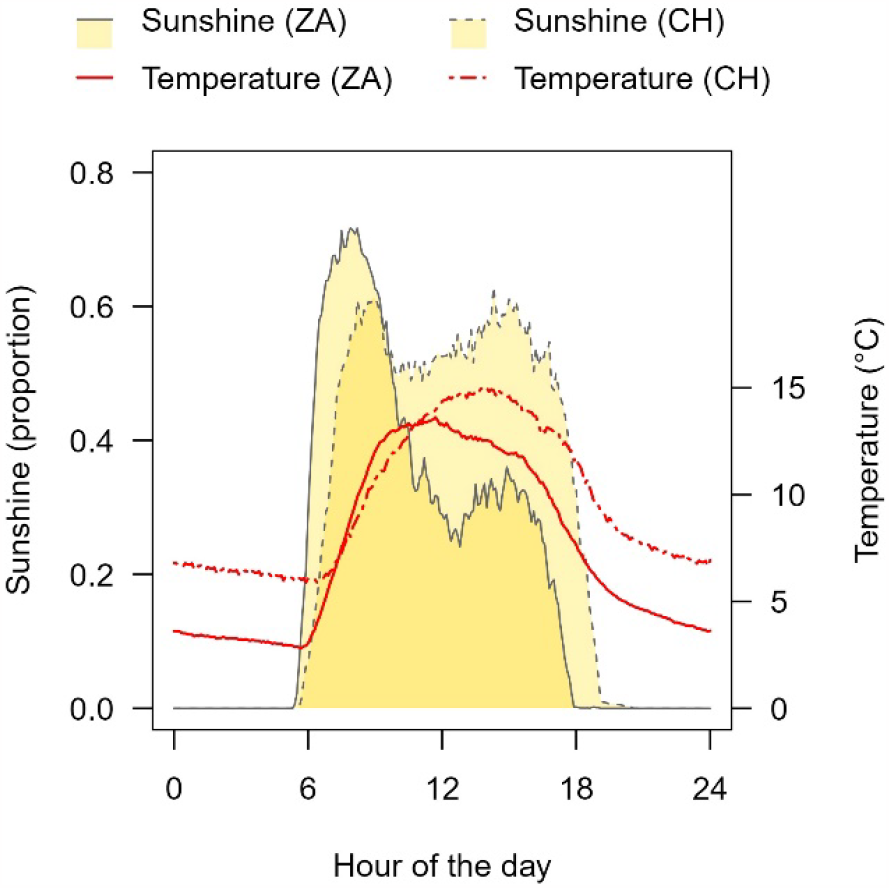
Average diel profiles of image-derived sunshine (yellow shading and grey lines) across high elevation sites in Switzerland (CH, dashed lines) and South Africa (ZA, solid lines) for 120 days following summer solstice. We also present mean air temperatures from on-board sensors on cameras, measuring air temperature every 5 minutes at 62cm above ground (red lines). Data are displayed with a resolution of 0.1 hours (240 time slices per day). The ZA site is characterized by early morning sunshine and afternoon shade. Sunshine was classified using a deep learning ensemble trained using the full set of 8,953 labelled images.

### Effects of experimental warming on snowmelt

We found clear impacts of OTCs on the prevalence and duration of snow cover (Fig. 5). During the first (and longest) recorded snowfall, the snow melted completely after around 2.5 days in warmed plots. This contrasted with unwarmed plots, where snow persisted for up to four days (Fig. 5). Beyond temporal variation, we also captured fine-scale spatial variation in retention of snow cover – especially among warmed plots (Fig. 5, left). Sensor-derived temperatures showed evidence of warming within OTCs, especially during prolonged sunshine, which probably contributed to advanced snowmelt. However, snow was often less prevalent across warmed plots even immediately after snowfall (Fig. 5). This suggests that OTCs also intercept a fraction of falling snow, which could delay snowpack formation.

**Figure 5.**
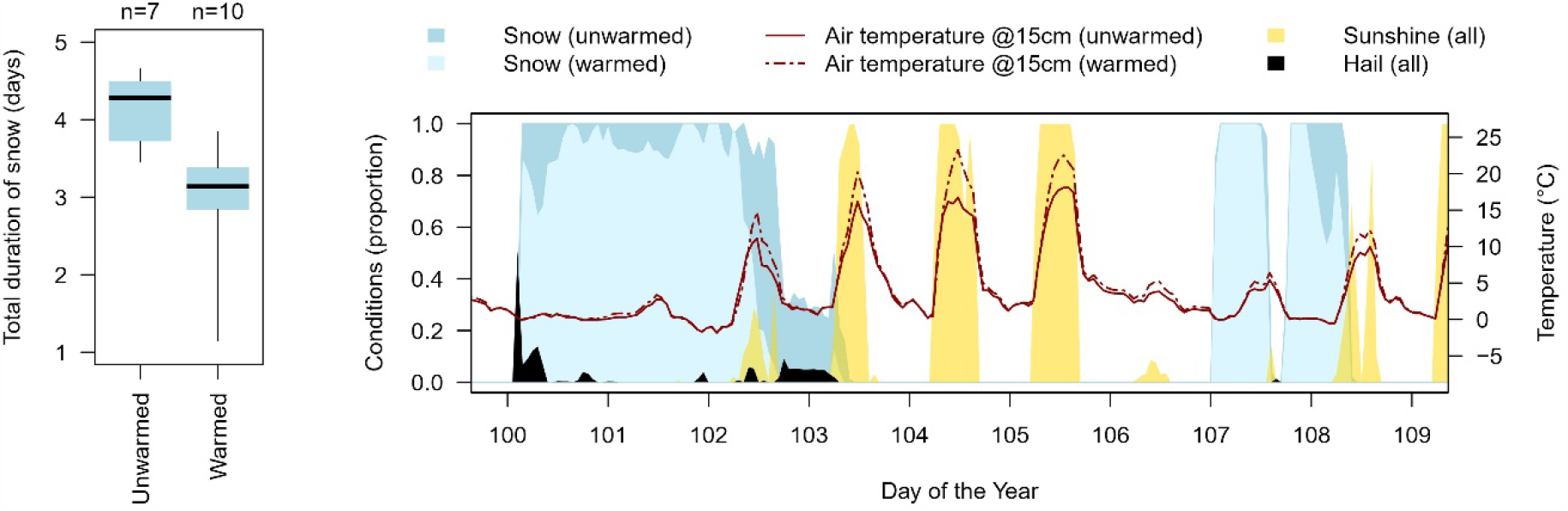
**(left)** Duration of snow, derived from images, across unwarmed plots and open-top chamber (OTC) warmed plots at the high elevation site in South Africa. Boxes capture the median and interquartile range, while whiskers capture the range of the data. Three cameras failed before the snows and are excluded here. **(right)** Proportion of snow over time across unwarmed (dark blue) and warmed (light blue) plots. Snow events were often shortened or dampened within OTCs. Furthermore, during periods of sunshine (yellow), temperature spikes recorded by TMS4 loggers were more intense in warmed plots (dashed red line) than unwarmed plots (solid red line). Small proportions of snow images were misclassified as hail (black), and this was often due to fog. Weather conditions were classified using a deep learning ensemble trained using the full set of 8,953 labelled images.

Few studies have reported effects of OTCs on snow cover in autumn. Several studies have reported effects on snow depth in winter and snowmelt in spring, although observations are often infrequent or anecdotal (Bokhorst et al., 2013; Wipf & Rixen, 2010). Heavy snows are known to accumulate in OTCs over winter, increasing snow depth and insulating plants and soil (Hollister et al., 2023; Rixen et al., 2022); the resulting winter warming of OTCs often exceeds their spring and summer warming effects (Aerts et al., 2004; Bjorkman et al., 2015). Such snow accumulation leads to unpredictable effects of OTCs on snowmelt; it may remain unchanged, be delayed by a week, or be advanced by several weeks (Aerts et al., 2004; Marion et al., 1997; Wipf & Rixen, 2010). Here we show that in early autumn, experimental warming can reduce the duration of snow cover – although this may partly result from interception of snowfall by OTCs. Furthermore, we demonstrate how to efficiently quantify persistence of snow at high spatiotemporal resolutions. This is a vital contribution given the impacts of snow cover on plant and soil ecosystem processes (Möhl et al., 2022), and the prevalence of OTCs in climate change research (Bjorkman et al., 2015; Hollister et al., 2023; Rixen et al., 2022).

### Effects of sunshine on bumblebee foraging

Sunshine is a vital factor affecting the activity of flower visiting insects, especially in alpine environments (Bergman et al., 1996). Our data demonstrate how sunshine drives fluctuations in ambient temperature at our site (Fig. 3), but also the cumulative daytime temperatures experienced by individual *T. pratense* inflorescences (Fig. 6, top left). We also observe a strong effect of sunshine exposure on the number of *Bombus* foraging visits to each inflorescence (Fig. 6, bottom left), probably partly mediated by ambient temperatures. However, we found that sunshine days marginally outperformed degree days when predicting the number of *Bombus* visits to an inflorescence (ΔAIC = -0.98). As such, we suggest that sunshine also has direct effects on *Bombus* foraging activity. For example, solar radiation can directly raise thoracic temperatures of bees, relieving a major constraint on flight behavior (Corbet et al., 1993).

**Figure 6.**
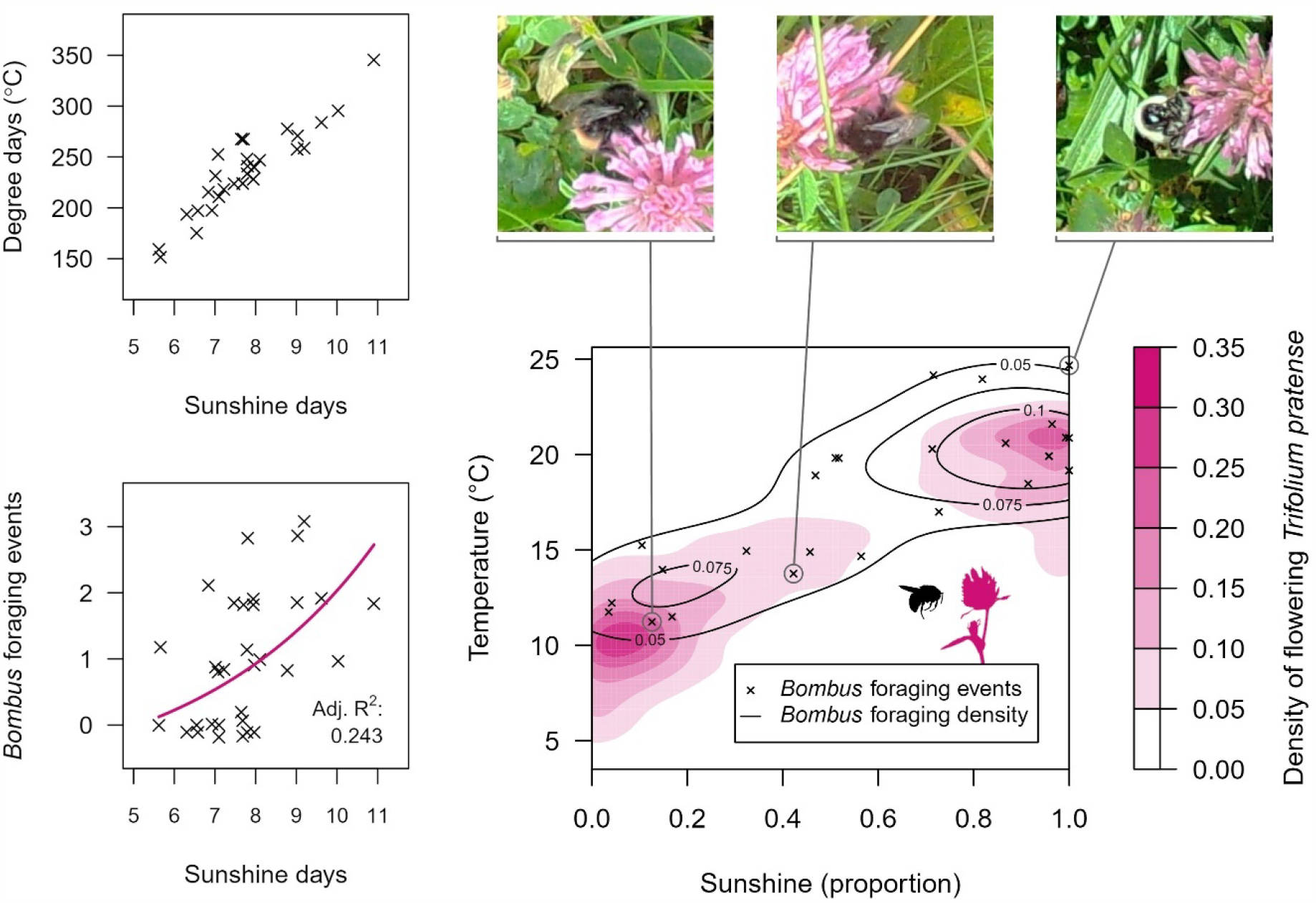
**(top left)** The cumulative daytime temperatures experienced by *Trifolium pratense* inflorescences, derived from temperature sensors, are highly positively correlated with sunshine exposure, derived from images. **(bottom left)** Sunshine exposure of an inflorescence increases the number of recorded *Bombus* foraging events. A linear model effectively predicts the natural log of the number of *Bombus* foraging events (pink line). **(right)** Sunshine and temperature preferences of foraging *Bombus* bees during late summer days at a high elevation site in Switzerland. The plot shows the temporal niche overlap between flowering *Trifolium pratense* (pink) and their *Bombus* bee pollinators (black crosses and lines) along axes of image-derived sunshine and sensor-derived temperature. The density of both flowering *T. pratense* and *Bombus* visits are bimodal with respect to sunshine and temperature, but *Bombus* bees are more likely to visit during warm and sunny periods. Image crops corresponding to three *Bombus* foraging events are shown above the plot.

We overlaid the density of *Bombus* foraging events with the density of flowering *T. pratense* on a two-dimensional surface of sunshine and temperature. Even though *Bombus* visits are contingent on the presence of flowering *T. pratense*, we found clear evidence of microclimatic and micrometeorological mismatch. *Trifolium pratense* inflorescences were most available at ∼10°C with little or no sunshine, with a secondary peak at ∼21°C during constant sunshine (Fig. 6, right). However, *Bombus* foraging events were completely absent below 10°C, and disproportionately prevalent during warm temperatures and intermediate sunshine. Camera-based monitoring, with automated extraction of meteorological by-catch, allows us to quantify the constraints and preferences of species at unprecedented spatial and temporal resolutions.

### Applications and future development

Micrometeorological variables extracted from images are highly complementary to those recorded by affordable microclimatic sensors. Unlike temperature and moisture, solar radiation is expensive and difficult to measure at high temporal resolution (Roales et al., 2013). Furthermore, cameras are perhaps the only in-field sensors that can record the prevalence of snow (Caparó Bellido & Rundquist, 2021) and hail at fine spatiotemporal resolutions. Crucially, the impacts of sunshine, snow, and hail on the abundance and phenology of wild species are significant, but rarely explored and poorly understood (Fernandes et al., 2011; Möhl et al., 2020, 2022; Valladares et al., 2016). Furthermore, cameras can capture variation in micrometeorology and organismal activity at very small spatial scales. This creates opportunities to study fine-scale microclimatic refugia, such as areas protected from sunshine or snow, which may be vital for species persistence in extreme environments (von Oppen et al., 2022). Cameras also record continuously at high temporal resolution, allowing the analysis of phenological mismatches not only across days of the year, but hours of the day (Alison et al., 2022). Finally, cameras generate biological and meteorological data that are tethered in both space and time. This is a particularly useful property to explore behavioral adaptations to micrometeorological conditions.

Previous studies have automated the extraction of snow-covered regions of phenocam images (Caparó Bellido & Rundquist, 2021; Julitta et al., 2014). Our approach of classifying entire images is simpler, and thus broader in application. Our model rapidly detects images with snow, but also those with sunshine or hail. Unlike previous models, ours is shown to perform well on images from novel camera placements and even a distant site in Svalbard. Above all, we present a dataset and methods to train future models that will be even more transferable. Future work should focus on the assembly of larger training datasets, representing a greater diversity of backgrounds, conditions, and wildlife camera domains. There is potential for deep learning models to extract hydrological information from images, for example rainfall or even rare cryogenic processes (Grab et al., 2021). Furthermore, models could be trained to extract meteorological by-catch from phenocams for vegetation (Brown et al., 2016) or camera traps for large animals (Hofmeester et al., 2020). A particularly exciting prospect would be to extract latent meteorological information from existing wildlife camera network datasets, containing millions of labelled organisms (Norouzzadeh et al., 2018). This could provide new insights into the occurrence and phenology of a huge variety of animal and plant species worldwide.

## Acknowledgements

This research was funded through the 2019–2020 BiodivERsA joint call for research proposals, under the BiodivClim ERA-Net COFUND programme, with the funding organizations Innovation Fund Denmark (grant no. 0156-00022B), the Department of Science and Innovation Republic of South Africa (grant no. DSI/CON 0000/2021), the Research Council of Norway, the Swiss National Science Foundation (grant no. 20BD21_193809), the Swedish Research Council for Environment, Agricultural Sciences and Spatial Planning and the German Research Foundation.

