## Supplementary information for "Deep learning to extract the meteorological by-catch of wildlife cameras"

**Table S1.** Rare meteorological events were sorted into three tiers based on duration and representation in the systematic subset. This determines the sampling protocol for strategic sampling and allows for more intensive sampling of the rarest events.

| Event duration | Representation | Strategic sampling protocol | Applies to: |
|---|---|---|---|
| > 1 day | < 100 images | Extract one image per-camera per-hour during the two non-consecutive days with highest prevalence of the given condition | Snow at ZA high site<br>Snow at CH high site<br>Frost at CH high site |
| < 1 day | < 100 images | Extract one image per-camera per-hour during three non-consecutive 12h windows with highest prevalence of the given condition | Hail at ZA high site |
| < 2 hours | < 10 images | All images across all cameras in the 2h window with the highest prevalence of the given condition. | Hail at CH high site |

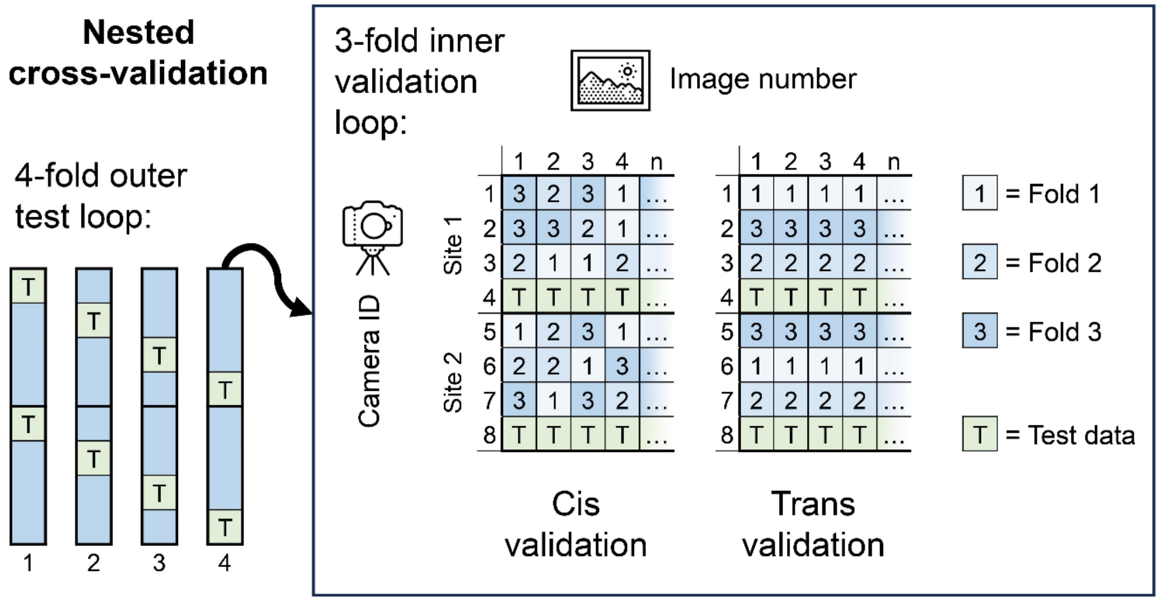


**Figure S1.** A simplified example of the nested cross-validation procedure – note, our study uses inner and outer validation loops with five and six folds respectively. Here, the outer test loop determines which of *k*=4 folds of cameras will be held back as out-of-sample test data. For each iteration of the outer test loop, an inner validation loop determines which of *k*-1=3 folds of images are used for early stopping. The delineation of folds in the inner validation loop is done using one of two methods: “Cis” or “Trans”. Cis validation involves naïve random splitting of images from all cameras. Trans validation splits based on cameras rather than images, as in the outer test loop. During Trans validation, a given camera never contributes to both model training and validation. Data splitting is always stratified with respect to sampling site.

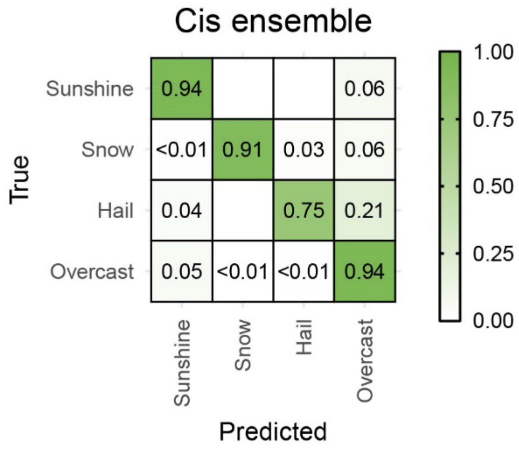


**Figure S2.** Confusion matrix of average predictions of an ensemble model following Cis validation (left) and Trans validation (right). Six Cis ensembles and six Trans ensembles were used to classify images from distinct hold-out test cameras.

**Appendix S1. Further details on image labelling.** We initially labelled evidence of sunshine, hail, snow and frost. Sunshine was evidenced by clear shadows cast by vegetation; hail by pellets of ice and ensuing slush; snow by snowflakes and snow cover, as well as ensuing slush; and frost by ice crystals forming on vegetation. Frost differed between the two regions; in ZA, heavy white frost erupted from the vegetation to form snowy crystals. This strongly resembled, and was often accompanied by, precipitated snow. In CH, frost was light and difficult to separate from dew. As such, frost was included in the snow class in ZA but the overcast class in CH. Overcast images did not adhere to a strict meteorological definition of overcast, being defined by the absence of sunshine, snow or hail. They comprised images of dry or wet vegetation, day and night, with occasional fog or light frost. A few images containing both sunshine and snow were included in the snow class.

**Appendix S2. A primer on CNN parameters and concepts.** During training of a CNN, training images are fed to an algorithm iteratively in computationally manageable **batches** of a given number of images. Model parameters are updated after each iteration, based on a given **optimizer** and a **learning rate**, to minimize a **loss function** (effectively a measure of how wrong the model's predictions are for a given batch of images). An **epoch** is a full cycle of iterations that spans the entire training dataset. The batch size, the optimizer, the learning rate, and the number of epochs are hyperparameters. **Overfitting** means an increase in model performance with respect to the training dataset (measured as the training loss), but a decrease in model performance with respect to independent observations in a validation dataset (measured as the validation loss). **Early stopping** means finding the epoch when training should be stopped to maximize model performance with respect to a validation dataset.

**Appendix S3. Further details on Trans validation and crogging ensembles.** Several obstacles hinder deep learning approaches to extract biotic or abiotic data from images. Chief among these is model transferability to novel camera deployments or locations, which often goes untested. This issue is exacerbated by redundancy or shared information between images from the same time-series; training datasets with tens of thousands of images may represent <100 individual deployments (Bjerge et al., 2023). To prevent overfitting to specific image backgrounds, studies must deploy many cameras across distinct locations. Furthermore, the presence of shared information between training, validation and test datasets can cause a phenomenon known as "information leakage", leading to underestimation of prediction error (Wieczorek et al., 2022). This can be avoided by splitting images appropriately between training, validation, and test datasets (Beery et al., 2018, 2020). In this study, we compared two data splitting methods for validation in our study: "Cis" and "Trans" (Fig. S1). The Trans validation approach withholds entire deployments for an especially rigorous validation process. Few studies applying deep learning to wildlife camera imagery have used a Trans data splitting process for model validation, though some have used it for model testing (Beery et al., 2018, 2020).

A major limitation of the Trans validation approach is that it greatly reduces the effective size of the training dataset, especially if the number of deployments is far smaller than the number of images. In the field of forecasting, an approach called cross-validation aggregation (crogging) has helped to overcome this barrier by allowing distinct time-series to contribute to both training and validation datasets – but not in the same model (Barrow & Crone, 2013). Hitherto untested for wildlife camera applications, crogging is an ensemble modelling approach that combines the benefits of cross-validation and model aggregation. Crogging can outperform other ensemble methods such as bootstrap aggregation (bagging; Barrow & Crone, 2013), and has proven useful where there is non-independence in the data – for example when classifying multiple brain tissue images per patient (Beleites & Salzer, 2008) or forecasting from autocorrelated time-series (Barrow & Crone, 2016). However, there is the caveat that the number of folds used during crogging affects the proportion of data used for training vs. improving out-of-sample performance of each member model (Barrow & Crone, 2016; Krogh & Vedelsby, 1994).

We compared crogging ensembles using the Cis validation approach, which permits images from the same camera to be used in both training and validation sets, against those using the Trans validation approach, which does not (Beery et al., 2018, 2020). We found that performance of ensemble models from the Trans validation approach was comparable to that of the Cis validation approach, even though member models from the Trans validation approach saw 20% fewer cameras during training. Furthermore, our crogging ensemble models tended to outperform full models, which were trained on all available data for up to 29 epochs, retaining the model from the epoch with the lowest training loss. In any case, the average difference between member and ensemble models was generally greater when using the Trans validation approach (Table 1). Our results highlight that the combination of crogging and Trans validation approaches demands further exploration in deep learning applications to images from wildlife cameras.

Barrow, D. K., & Crone, S. F. (2013). Crogging (cross-validation aggregation) for forecasting—A novel algorithm of neural network ensembles on time series subsamples. *Proceedings of the International Joint Conference on Neural Networks*. https://doi.org/10.1109/IJCNN.2013.6706740

Barrow, D. K., & Crone, S. F. (2016). Cross-validation aggregation for combining autoregressive neural network forecasts. *International Journal of Forecasting*, *32*(4), 1120–1137. https://doi.org/10.1016/j.ijforecast.2015.12.011

Beery, S., Van Horn, G., & Perona, P. (2018). Recognition in Terra Incognita. *Lecture Notes in Computer Science (Including Subseries Lecture Notes in Artificial Intelligence and Lecture Notes in Bioinformatics)*, *11220 LNCS*, 472–489. https://doi.org/10.1007/978-3-030-01270-0_28

Beery, S., Liu, Y., Morris, D., Piavis, J., Kapoor, A., Meister, M., Joshi, N., & Perona, P. (2020). Synthetic examples improve generalization for rare classes. *Proceedings - 2020 IEEE Winter Conference on Applications of Computer Vision, WACV 2020*, 852–862. https://doi.org/10.1109/WACV45572.2020.9093570

Beleites, C., & Salzer, R. (2008). Assessing and improving the stability of chemometric models in small sample size situations. *Analytical and Bioanalytical Chemistry*, *390*(5), 1261–1271. https://doi.org/10.1007/s00216-007-1818-6

Bjerge, K., Alison, J., Dyrmann, M., Frigaard, C. E., Mann, H. M. R., & Høye, T. (2023). Accurate detection and identification of insects from camera trap images with deep learning. *PLOS Sustainability and Transformation*, *2*(3), e0000051. https://doi.org/10.1371/journal.pstr.0000051

Krogh, A., & Vedelsby, J. (1994). Neural Network Ensembles, Cross Validation, and Active Learning. *Advances in Neural Information Processing Systems*, *7*, 231–238.

Wieczorek, J., Guerin, C., & McMahon, T. (2022). K-fold cross-validation for complex sample surveys. Stat, 11(1), e454. https://doi.org/10.1002/sta4.454
